# Effect of tempol on post-thaw semen parameters and fertility during chicken semen cryopreservation

**DOI:** 10.1101/2022.03.31.486544

**Authors:** M. Shanmugam, R.K. Mahapatra

## Abstract

The present study evaluated addition of tempol during chicken semen cryopreservation on post-thaw semen parameters and fertility. Adult PD-1 line semen was cryopreserved using 4% dimethyl sulfoxide (DMSO) in Sasaki diluent (SD). In the semen cryomixture tempol (1 and 5 mM) was added at final concentrations. The semen with additives were filled in 0.5 ml French straws and exposed to liquid nitrogen vapours for 30 min and then stored in liquid nitrogen. The semen straw was thawed at 5°C for 100 sec and evaluated for sperm motility, live, abnormal and acrosome intact sperm. The seminal plasma was evaluated for lipid peroxidation. Fertilizing potential of the cryopreserved sperm was evaluated after insemination in the PD-1 line hens. The post-thaw sperm parameters were significantly (P<0.05) lower in the cryopreserved groups. The lipid peroxidation was significantly (P<0.05) higher in cryopreserved groups. The fertility was significantly (P<0.05) lower in all the cryopreserved groups. In conclusion, addition of tempol to cryopreservation mixture did not improve the post-thaw semen parameters or fertility.

---

Cryopreservation of chicken semen is a low-cost management tool for conservation. The desired fertility results in chicken similar to that obtained in cattle for practical use is still elusive due to various reasons such as line or breed variability (Long 2006). The concentration of polyunsaturated fatty acids in chicken sperm is high and is prone for lipid peroxidation (Surai et al. 2001). The limited antioxidant system present in semen is overwhelmed during cryopreservation process where high levels of reactive oxygen species are formed resulting in higher lipid peroxidation damage to the sperm membrane (Partyka et al. 2012). Therefore, addition of antioxidants in the semen cryodiluent will help in reducing the damage occurring during freezing and thawing process.

Tempol (4-hydroxy2, 2, 6, 6-tetramethylpiperidine-1-oxyl) is a low molecular weight cyclic nitroxide compound that has superoxide dismutase (SOD) enzyme mimetic activity. It has very good cell permeability and has been used in human and alpaca semen cryopreservation (Santiani et al. 2013; Bateni et al. 2014; Azadi et al. 2017). To our knowledge there is no report on the use of tempol during chicken semen cryopreservation. Therefore, the present study aimed to evaluate addition of tempol in cryopreservation mixture and determine its effect on post-thaw semen quality and fertility in chicken.

The experimental procedure was carried at the institute poultry farm. Adult PD-1 birds were housed individually in cages in an open-sided house. Feed and water were available *ad libitum* throughout the experimental period. The experimental procedures were approved by the Institutional Animal Ethics Committee.

PD-1 roosters of 40 weeks age were used for semen collection by dorso-abdominal massage method (Burrows and Quinn 1937). The collected semen was pooled and cryopreserved. An aliquot of pooled fresh semen was used for estimating sperm motility, live and abnormal sperm, and acrosome intact sperm. The semen was centrifuged at 3000 x g for 5 min to separate seminal plasma that was stored until analysis. Sasaki diluent (D (+)-glucose-0.2 g, D (+)-trehalose dehydrate-3.8 g, L-glutamic acid, monosodium salt-1.2 g, Potassium acetate-0.3 g, Magnesium acetate tetrahydrate-0.08 g, Potassium citrate monohydrate-0.05 g, BES-0.4 g, Bis-Tris-0.4 g in 100 ml distilled water, final pH 6.8; Sasaki et al. 2010) was used for cryopreserving the semen. The semen for cryopreservation was kept at 5°C for 30 min and then diluted in equal volume of diluent containing 8% DMSO. Tempol (CAS no.2226-96-2; Sigma-Aldrich Co., St. Louis, USA) was added to this mixture at 1 and 5 mM final concentration. The semen cryomixture was loaded in 0.5 ml French straws and placed 4.5 cm above liquid nitrogen exposing it to nitrogen vapours for 30 min after which they were stored in liquid nitrogen until further use. The semen cryopreservation and post-thaw evaluation was done on six occasions for progressive sperm motility, live sperm, abnormal sperm, and intact sperm acrosome. After storing for a minimum period of seven days the plastic straws were thawed at 5°C for 100 sec. Thawed semen was inseminated into 41 weeks old PD-1 hens (12 hens/treatment) with 200 million sperm. The insemination was repeated three times at four days interval. The fresh semen inseminated group served as control. After insemination the eggs were collected and incubated under standard incubation conditions. Candling of eggs was done on 18^th^ day of incubation from which fertility data was obtained.

The progressively motile sperm were scored subjectively after placing a drop of semen on a Makler chamber and examining under 20x magnification.

The live and abnormal sperm were assessed using Eosin-Nigrosin stain (Campbell et al. 1953). A semen smear was prepared after mixing a drop of semen and a drop of Eosin-Nigrosin stain, air dried and examined under high power (1000x) magnification. The live membrane intact sperm that were clear in appearance were counted and percent live sperm calculated. A total of 200 sperm were counted in each slide. The abnormal sperm percent assessed based on morphological abnormalities were also estimated in the same slides.

The sperm acrosome intactness was estimated as per the earlier reported protocol (Pope et al. 1991). Semen (10 μl) was mixed with equal volume of stain (1% (wt/vol) rose Bengal, 1% (wt/vol) fast green FCF and 40% ethanol in citric acid (0.1 M) disodium phosphate (0.2 M) buffer (McIlvaine’s, pH 7.2-7.3) and left for 70 sec. On a glass slide a smear from the mixture was made, dried and evaluated at high magnification (1000x). Sperm having intact acrosome was identified by the blue stained acrosomal caps while no stained cap could be observed in the acrosome reacted sperm. The acrosome intact sperm percent was calculated by counting a minimum of 200 sperm in each sample.

Lipid peroxidation in seminal plasma was measured by the thiobarbituric acid method (Hsieh et al. 2006). Semen samples were centrifuged at 3000 x g for 5 min to separate seminal plasma and was stored until analysis. In each test tube 0.9 ml of distilled water and

0.1 ml seminal plasma was added and mixed. This was followed by addition of 0.5 ml of thiobarbituric acid reagent. Tubes without sample was processed that acted as blank. The tubes were kept in boiling water bath for one hour. After cooling the tubes absorbance was measured against blank at 534 nm using spectrophotometer.

All the data were analyzed in SAS 9.2 and P<0.05 was considered significant. The fresh and cryopreservation treatments were compared by one-way ANOVA with Tukey’s post hoc test. The percent value data were arcsine transformed and then analysed.

All the semen parameters studied and fertility were significantly (P<0.05) reduced after cryopreservation (Table 1). Seminal plasma lipid peroxidation was significantly (P<0.05) higher in cryopreserved groups. Addition of tempol during cryopreservation had no significant (P>0.05) effect on improving post-thaw sperm motility, seminal plasma lipid peroxidation and fertility.

**Table 1:** Effect of inclusion of Tempol during chicken semen cryopreservation.

| Parameters | Fresh semen | 4% DMSO | 4% DMSO +<br>Tempol 1 mM | 4% DMSO +<br>Tempol 5 mM |
| --- | --- | --- | --- | --- |
| Progressive sperm motility (%) | 65.0 ± 1.8<sup>a</sup> | 21.67 ± 1.67<sup>b</sup> | 21.67 ± 1.67<sup>b</sup> | 23.33 ± 1.05<sup>b</sup> |
| Live sperm (%) | 77.58 ± 2.7<sup>a</sup> | 30.98 ± 1.86<sup>b</sup> | 28.58 ± 1.17<sup>b</sup> | 30.15 ± 2.26<sup>b</sup> |
| Abnormal sperm (%) | 1.8 ± 0.29 | 1.8 ± 0.18 | 1.8 ± 0.27 | 2.0 ± 0.14 |
| Acrosome intact sperm (%) | 93.0 ± 0.93<sup>a</sup> | 79.83 ± 4.39<sup>ab</sup> | 80.17 ± 3.48<sup>ab</sup> | 76.13 ± 4.48<sup>b</sup> |
| Seminal plasma lipid peroxidation (nM MDA/ml) | 1.97 ± 0.32<sup>b</sup> | 6.75 ± 1.23<sup>a</sup> | 4.41 ± 0.34<sup>ab</sup> | 5.03 ± 0.53<sup>a</sup> |
| Fertility (%) | 91.6 ± 2.47<sup>a</sup> | 16.33 ± 7.57<sup>b</sup> | 0<sup>b</sup> | 6.67 ± 6.67<sup>b</sup> |
| Number of eggs incubated | 75 | 55 | 83 | 44 |

Values are Mean±SE.
Values having different superscripts in a row differ significantly (P<0.05).

The harmful effects of semen cryopreservation could be observed in the present study in terms of higher seminal plasma lipid peroxidation level. Addition of tempol could not reduce this higher lipid peroxidation level. Tempol a six-membered cyclic nitroxide and SOD mimetic rapidly reacts with superoxide anions and prevents the progression of Fenton reaction (Samuni et al. 1990). In the present study if tempol had neutralized superoxide anion the level of lipid peroxidation in the treatments where it was added should have reduced, however, this did not occur. Tempol used at 1mM concentration during alpaca semen cryopreservation had been shown to improve post-thaw sperm motility, functional sperm membrane integrity and reduce sperm DNA fragmentation (Santiani et al. 2013). A low dose of 5μM tempol improved sperm motility and viability while reducing the sperm DNA fragmentation (Bateni et al. 2014; Azadi et al. 2017). During ram liquid semen cooled storage tempol at 2mM improved sperm motility and fertilization rate (Mara et al. 2005). Thus, the results of use of tempol in semen storage indicates its ability to protect sperm cell during preservation. In the present study the levels of tempol used in the cryopreservation medium might have not produced any beneficial effects on post-thaw semen parameters. Future studies should be carried out with tempol using different levels during chicken semen cryopreservation to observe any beneficial effects.

It is concluded that addition of tempol at 1 and 5 mM concentrations during chicken semen cryopreservation had no effect on post-thaw semen parameters or fertility.

